# Modulation of Transient receptor potential melastatin 3 by protons through its intracellular binding sites

**DOI:** 10.1101/2020.10.08.331454

**Authors:** Md Zubayer Hossain Saad, Liuruimin Xiang, Yan-Shin Liao, Leah R. Reznikov, Jianyang Du

## Abstract

Transient receptor potential melastatin 3 channel (TRPM3) is a calcium-permeable nonselective cation channel that plays an important role in modulating glucose homeostasis in the pancreatic beta cells. However, how TRPM3 is regulated under physiological and pathological conditions is poorly understood. In this study, we found that both intracellular and extracellular protons block TRPM3 through its intracellular binding sites. We demonstrated that external protons indirectly block TRPM3, whereas internal protons inhibit TRPM3 directly with an inhibitory pH_50_ of 6.9 ± 0.11. We identified three titratable residues, D1059, D1062, and D1073, at the inner vestibule of the channel pore that contribute to pH sensitivity. The mutation of D1073Q reduces TRPM3 current intensity and pH sensitivity; Replacement of Asp 1073 by Gln 1073 changes the reduction of TRPM3 outward current by low external pH 5.5, from 62 ± 3 % in WT to 25 ± 6.0 % in D1073Q. These results indicate that D1073 is not only essential for intracellular pH sensitivity, but it is also crucial for TRPM3 channel gating. In addition, a single mutation of D1059 or D1062 enhances pH sensitivity. In summary, our findings provide a novel molecular determinant for pH regulation of TRPM3. The inhibition of TRPM3 by protons may indicate an endogenous mechanism governing TRPM3 gating and its physiological/ pathological functions.

## Introduction

Transient receptor potential channels (TRP channels) are membrane proteins that facilitate the interpretation of external stimuli, and allow organisms to readily sense the environment (Clapham 2003). These stimuli include temperature (Vandewauw et al. 2018; Brauchi and Orio 2011), voltage (Clapham 2003; Brauchi and Orio 2011; Montell et al. 1985), mechanical force (osmolarity (Strotmann et al. 2000; Liedtke and Friedman 2003; Quallo et al. 2015), pressure (Suzuki et al. 2003), stretch (Strotmann et al. 2000; Hardie and Franze 2012; Maroto et al. 2005), gravity (Sun et al. 2009)), light (Montell et al. 1985; Minke 1977; Hardie 2014), proton concentration (Gerdes et al. 2007; Chandrashekar et al. 2006; Semtner et al. 2007), and various chemical signals (Caterina et al. 1997; McKemy, Neuhausser, and Julius 2002; Everaerts et al. 2011). TRP channels form homomeric or heteromeric cation channels that can be selective or non-selective to cations; and most of the TRP members are permeable to Ca^2+^ (Pan, Yang, and Reinach 2011). Upon activation, TRP channels change membrane potential or intracellular calcium concentration ([Ca^2+^]_i_) to promote downstream signal transductions. Depending on sequence homology and channel architecture, TRP channels are divided into seven subfamilies, namely, TRPA (Ankyrin), TRPC (Canonical), TRPM (Melastatin), TRPML (Mucolipin), TRPP (Polycystin), TRPV (Vanilloid), and TRPN (NompC, no vertebrate member). 27 vertebrate members of these subfamilies are expressed in humans (Venkatachalam and Montell 2007; Uchida et al. 2019).

TRPM3 belongs to the subfamily of TRPM (Melastatin). TRPM3 is a non-selective cation channel that is permeable to Ca^2+^, Na^+^, Mn^2+^ and Mg^2+^ ions with a permeability ratio of P_Ca_/P_Na_ of 1.57 ± 0.31 (Grimm et al. 2003). RT-qPCR analyses have shown expression of human (hTRPM3), mouse (mTRPM3) and rat (rTRPM3) TRPM3 in a variety of tissues, with the most abundant expression in the brain, kidney, pituitary gland and adipose tissues (Oberwinkler and Philipp 2014; Zamudio-Bulcock et al. 2011). In sensory neurons, TRPM3 functions as a noxious heat sensor; TRPM3 deficient mice lack the normal response to noxious heat and do not develop inflammatory heat hyperalgesia (Vandewauw et al. 2018; Vriens et al. 2011). Vangeel et al. recently demonstrated functional expression of TRPM3 in human sensory neurons; where a large subset of nociceptor neurons express TRPM3, and TRPM3 has been suggested to be a potential drug target for novel analgesics (Vangeel et al. 2020; Moran and Szallasi 2018). Although, role of TRPM3 in central nervous system has not been explored in detail yet, a recent study has shown high-level expression of TRPM3 in mouse CA2 and CA3 hippocampal neurons and in the dentate gyrus. Field potential recordings also showed that TRPM3 agonists inhibit synaptic transmission and plasticity, and reduce long-term potentiation (LTP) in mouse hippocampus (Held et al. 2020). Moreover, human genetic analyses have linked TRPM3 mutations with intellectual disability, epilepsy, inherited cataract and glaucoma (Dyment et al. 2019; Bennett et al. 2014). In cardiovascular system, TRPM3 localizes in the perivascular nerves of mouse mesenteric arteries, and induces vasodilation by stimulating CGRP (calcitonin gene-related peptide) receptors (Alonso-Carbajo et al. 2019). TRPM3 forms homomultimeric channels, which are constitutively active (Grimm et al. 2003). In addition, TRPM3 channel can also be activated by endogenous neurosteroid pregnenolone sulfate (PS), nifedipine, and clotrimazole. Mefenamic acid, diclofenac, progesterone and favanones have been reported to inhibit TRPM3 (Wagner et al. 2008; Uchida et al. 2019). PS has been shown to increase neurotransmitter release, strengthen synaptic transmission and modulate synaptic plasticity (Smith, Gibbs, and Farb 2014). However, whether TRPM3 contribute to any of these neuronal functions of PS or not, is not clear (Zamudio-Bulcock et al. 2011). One study has shown that PS-induced potentiation of spontaneous glutamate release in Purkinje neurons of developing rats is mediated by TRPM3 (Zamudio-Bulcock et al. 2011). In non-neuronal cells, PS upregulates activator protein 1 (AP-1) and early growth response protein 1 (Egr-1) transcriptional activity, which can be blocked by TRPM3 antagonists (Lesch, Rubil, and Thiel 2014). TRPM3 also upregulates c-Jun and c-Fos promoter activity and stimulates CRE-controlled reporter gene transcription in insulinoma and pancreatic β-cells, in a TRPM3-dependent manner (Muller, Rossler, and Thiel 2011). In vascular smooth muscle cells, PS increases [Ca^2+^]_i_ and modulates contractile responses, which can be inhibited by TRPM3 inhibitors (Naylor et al. 2010). PS also increases [Ca^2+^]_i_ in fibroblast-like synoviocytes and suppresses the secretion of hyaluronan via TRPM3 (Ciurtin et al. 2010).

Extracellular and intracellular protons modulate ion channel activity. For example, extracellular acidification activates acid-sensing ion channels (ASICs), G-protein coupled inward rectifier K^+^ channels, nifedipine sensitive L-type Ca^2+^ channels, and acid-sensitive Cl^-^ channels; while inhibiting two-pore domain K^+^ channels (TASK-1, TASK-2, TASK-3, TALK-1, and TALK-2), and reducing potency of ionotropic purinoceptors - P2X1, P2X3, P2X4 and P2X7. Intracellular acidosis activates inward rectifier K^+^ channel family protein Kir6.1, K2p channel TREK-1 and TREK-2; however, Kir1.1, Kir4.1, and Kir5.1 activity is inhibited by both intracellular and extracellular decreases in pH. Gap junction channels, connexins, are also inhibited by intracellular acidosis. In addition, both intracellular and extracellular acidification block the K2p channel TRESK, as well as depress or inhibit TWIK-1 and TWIK-2. Some of the ion channels, such as, TASK-2, TRESK, TALK-1, and TALK-2, which are blocked at low pH can be activated or gated open by alkalization. Protons cannot activate or inhibit P2X2 and P2X5 homomultimers but decreases potency and efficacy of ATP gating of P2X5, and sensitizes P2X2 receptors to ATP. To make things even complex, protons can have a variable effect depending on the subunit composition of heteromeric ion channels (Holzer 2009). TRP channels are no exception regarding variable activity in response to pH. Specifically, TRPV1, TRPV4, and TRPC4 are activated by a reduction of pH. In contrast, TRPC5 currents are increased by an acidic pH until 6.0 is reached, at which point further decreases in pH reduce current (Holzer 2009). PKD2L1 (TRPP2) expressing neurons show action potentials in response to citric acid (Huang et al. 2006), whereas intracellular and extracellular pH inhibits TRPM2 (Du, Xie, and Yue 2009). Yet very little is known about how pH regulates TRPM3. Thus, to understand the role of pH on TRPM3 activity, we studied how TRPM3 activation by PS responds to different extracellular and intracellular pH conditions. In all experiments, we activated TRPM3 by external application of PS, and all subsequent mentions of TRPM3 activity in this manuscript must be considered as TRPM3 activity in response to PS, and not TRPM3 constitutive activity. As PS induced TRPM3 currents are almost two orders of magnitude higher than TRPM3 constitutive currents (Wagner et al. 2008; Grimm et al. 2003; Lee et al. 2003), we concluded that for our experiments, it is reasonable to exclude the effects of constitutive TRPM3 activity.

## Materials & Methods

### Plasmid and molecular biology

The cDNA of human TRPM3 channel (accession number AJ505026) with C-terminal GFP tag was provided by C. Harteneck (University of Tübingen, Tübingen, Germany) (Grimm et al. 2003). Alternative splicing patterns of TRPM3 is highly conserved across human and rodents (Oberwinkler et al. 2005). To date, 25 isoforms of mTRPM3 protein have been identified, including a recently discovered variant - TRPM3γ3. Splicing events affect exons 8, 13, 15, 17, 20, 24 and 28 of TRPM3. α variants lack exon 2, β variants lack exon 1, and γ variants lack a large part of exon 28 (Uchida et al. 2019; Oberwinkler and Philipp 2014). Our hTRPM3 cDNA contains all 30 exons, where 389 amino acids in exon 28 has been replaced with alternative carboxy terminus of 7 residues; this truncation does not affect any functional activity of the ion channel (Oberwinkler and Philipp 2014; Grimm et al. 2003). Mutations of hTRPM3-GFP were generated by site-directed mutagenesis (performed by GENEWIZ Inc). The predicted mutations were verified by sequencing analysis.

### Cell culture and overexpression of hTRPM3-GFP and the mutants in HEK-293 cells

Human embryonic kidney (HEK) 293 cells were used to transiently overexpress wild-type hTRPM3-GFP and its mutants. The cells were grown in DMEM/F12 medium (Fisher Scientific, catalog no. MT10090CV) supplemented with 10% bovine growth serum (HyClone, catalog no. SH30541.03), 100 U/ml penicillin / 100 mg/ml streptomycin (Fisher Scientific, catalog no. SV30010) at 37°C in a 5% CO_2_-controlled, humidity-controlled incubator. Lipofectamine 2000 (Thermo Fisher Scientific, catalog no. 18324012) was used for the transfection of TRPM3 into the cells in a 35-mm culture dish according to the manufacturer’s instructions. Successfully transfected cells were identified by their fused GFP when illuminated at 480 nm excitation wavelength. Electrophysiological recordings were conducted between 36- and 48-hours post-transfection.

### Electrophysiology

All patch-clamp experiments were performed at room temperature (20–22°C). TRPM3 whole-cell currents were recorded using an Axopatch 200B amplifier. Data were digitized at 10 kHz and digitally filtered offline at 5 kHz. Patch electrodes were pulled by Sutter P-97 micropipette puller and fire-polished to resistance of 3-5 MΩ when filled with internal solutions. Series resistance (Rs) was compensated up to 90% to reduce series resistance errors to <5 mV. Cells in which Rs was >8 MΩ were discarded (Du, Xie, and Yue 2009). For whole-cell current recording, ramp voltage stimuli (250 ms duration) were delivered at 1-second intervals and the ranging from −100 to +100 mV. The internal pipette solution for whole-cell current recordings contained (in mM): 115 Cs-methanesulfonate (CsSO_3_CH_3_), 8 NaCl, 10 Cs-EGTA, 5 Na_2_-ATP and 10 HEPES, with pH adjusted to 7.2 with CsOH. In high intracellular Ca^2+^ experiments, 0.93 mM CaCl_2_ was added to the above-mentioned intracellular solution and EGTA was reduced to 1 mM, resulting in 1 μM free intracellular Ca^2+^. MaxChelator (https://somapp.ucdmc.ucdavis.edu/pharmacology/bers/maxchelator/downloads.htm) software from the University of California, Davis was used to calculate free [Ca^2+^]_i_.

To avoid proton activated chloride currents conducted by endogenous anion channels of HEK-293 cells (Lambert and Oberwinkler 2005), NaCl in standard Tyrode solution was replaced with Na-glutamate for all whole-cell current recordings. This external solution contained (in mM): 145 Na-glutamate, 5 KCl, 2 CaCl_2_, 1 MgCl_2_, 10 HEPES, and 10 glucose, with pH adjusted to 7.4 with glutamic acid. Internal and external acidic pH solutions were prepared as described previously with slight modifications (Du, Xie, and Yue 2009). In brief, 10 mM HEPES used in the solutions at pH 7.4 and 7.0 was replaced by 10 mM MES for the solutions at pH ≤ 6.0. Bath solutions containing 1 mM to 60 mM NH_4_Cl were prepared by decreasing Na^+^ concentrations to 85 mM in the solution to keep the osmolarity constant, and osmolarity was adjusted to 300 ± 10 mOsm with mannitol. In experiments designed to test protons permeability of TRPM3, pipette solutions contained (in mM): 120 NMDG, 108 glutamic acid, 10 HEPES, 10 EGTA, with pH adjusted to 7.2 with NMDG. External solutions for proton permeability test contained (in mM): 145 NMDG, 10 HEPES and 10 Glucose; and pH was adjusted with glutamic acid. To prepare the pH 5.5 external solution for proton permeability, 10 mM HEPES was replaced with 10 mM MES. PS was dissolved in DMSO to prepare 100 mM stock solution, and adequate volume of stock PS solution was added to the external solution to achieve required concentration. All the chemicals used in electrophysiological experiments were from Sigma-Aldrich.

### Data analysis

Statistical data were analyzed using GraphPad Prism 8. Pooled data are presented as mean ± SEM. Concentration-response curves were fitted by an equation of the form: *E = Emax{1/[1+(IC_50_/C)^n^]}* where *E* is the effect at concentration *C*, *E_max_* is the maximal effect, IC_50_ is the concentration for half-maximal effect, and n is the Hill coefficient (Du, Xie, and Yue 2009). Concentration of proton required for half-maximal inhibition is denoted by IC_50_ (when H^+^ concentration is expressed as molar concentration) and pH_50_ (when H^+^ concentration is expressed by pH value). Statistical comparison of two groups was performed by unpaired Student’s t-test, p < 0.05 was considered statistically significant. Statistical comparison of three or more groups was performed by one-way ANOVA with Tukey’s post hoc multiple comparison.

## Results

### Extracellular and intracellular acidic pH inhibit TRPM3

We studied the effects of low pH on TRPM3 by overexpressing hTRPM3-GFP in HEK-293 cells and recording whole-cell currents in response to PS. We found that low extracellular pH inhibits TRPM3, in a reversible manner (Fig. 1, A - D). To avoid proton activated endogenous anion channel conducted chloride currents, external Cl^-^ was replaced by glutamate (See Methods). Inhibitory effect of low extracellular pH (pH_o_) was only observed below pH 6.0. Specifically, at pH_o_ 7.0 and 6.0, recorded TRPM3 currents were equivalent to pH_o_ 7.4 (p > 0.05 in both groups). At a pH_o_ below 6.0, acidic conditions exhibited significant inhibition of TRPM3 (Fig. 1). pH_o_ 5.5, caused ~ 60% reduction in TRPM3 whole-cell current induced by PS. Fitting these data in a non-linear regression curve did not result in a well-fitted curve; we did not observe a concentration-dependent effect of extracellular pH on TRPM3 activity. This might suggest an indirect inhibition of TRPM3 by low pH_o_. In addition, as shown in Fig. 1A, application of low extracellular pH (pH≤5.5) with PS produced an initial activation of TRPM3 before blocking it. This suggested that the onset of low pH_o_ inhibition is slower than the PS activation, supporting the hypothesis that the inhibition of TRPM3 by low pH_o_ is indirect. We thus hypothesized that protons block TRPM3 by permeating through TRPM3 and binding on a cytoplasmic site. To test this hypothesis, we first investigated the effects of intracellular low pH (pH_i_) on TRPM3. Whole-cell TRPM3 currents were recorded using the low-pH pipette solutions (see experimental procedures), while keeping extracellular pH constant at 7.4 (Fig. 2). Low pH_i_ markedly reduced TRPM3 current in a concentration-dependent manner with a pH_50_ value of 6.90 ± 0.11 (outward current at +100mV) (Fig. 2C) and pH_50_ value of 6.90 ± 0.15 (inward current at −100mV) (Fig. 2D). There was no significant difference in the steady-state inhibition between inward and outward currents, concluding that there are no voltage-dependent effects of acidic pH_i_ on inward and outward TRPM3 currents (Fig. 2, C and D). To investigate the extent of modulation of TRPM3 by protons, we also introduced higher pH_i_ than the physiological pH_i_ of 7.2. Recorded TRPM3 current plateaued at about pH_i_ 7.6. Protons had similar inhibitory effects on both outward and inward TRPM3 currents (Fig. 2, A and B). Combined, our findings that pH_o_ 6.0 did not affect TRPM3 current but low pH_i_ had an inhibitory pH_50_ value of 6.9, suggested a higher pH sensitivity of TRPM3 in the cytoplasmic side.

**Figure 1.**
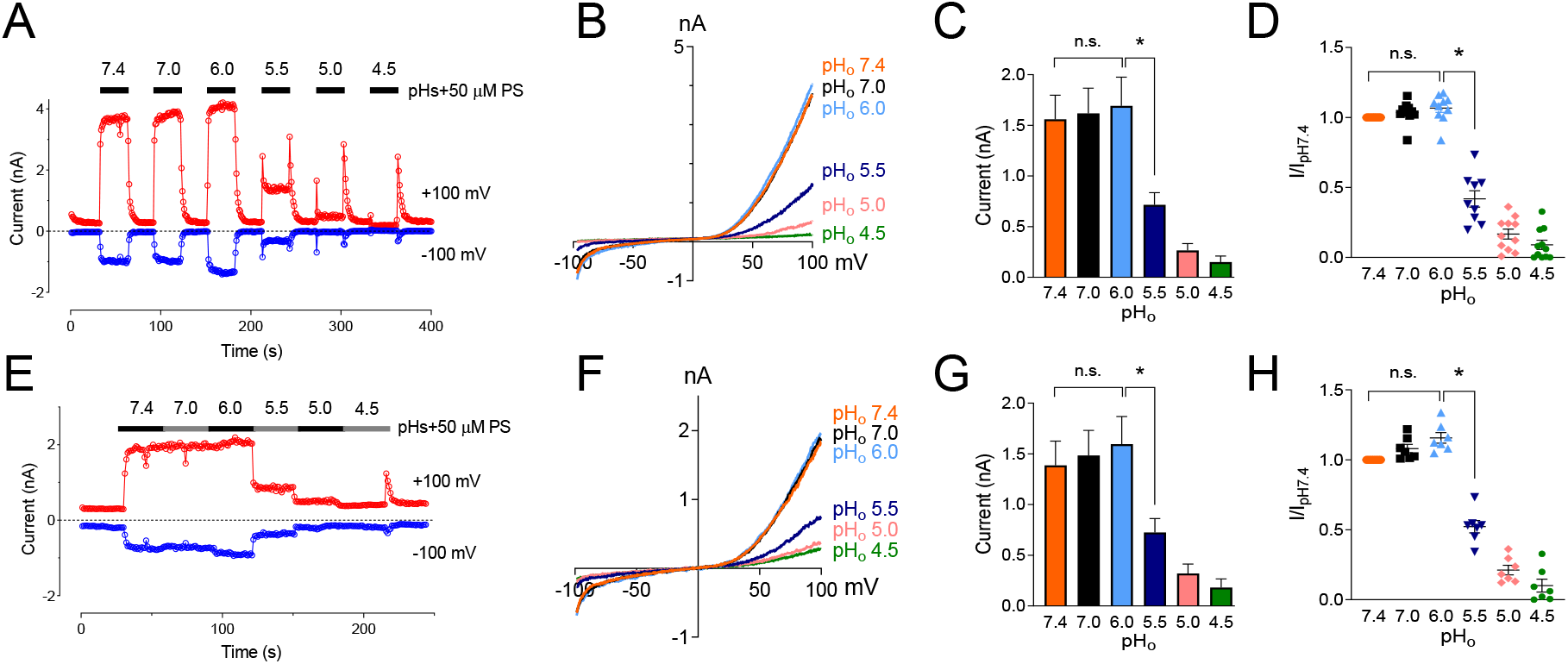
Inhibitory effect of extracellular acidic pH on TRPM3 activation by PS. (a) Time course of TRPM3 currents elicited by voltage ramps ranging −100 to +100 mV. Both inward and outward currents were completely and reversibly inhibited by pH_o_ 4.5. 50 uM PS was applied extracellularly and was washed with PS-free extracellular solution between subsequent PS application. Inward and outward currents were measured at −100 and +100 mV, respectively. (b) Representative recording of TRPM3 current in (a) by ramp protocols ranging from −100 to +100 mV at the indicated pH_o_. (c) Mean current amplitude of TRPM3 at the indicated pH_o_ in (a) (mean ± SEM; n = 11, * indicates p<0.05 by unpaired Student’s t-test; “n.s.” indicates not statistically significant). (d) Current amplitude at the indicated pH_o_ normalized to the current amplitude at pH_o_ 7.4 in (a). When compared with pH_o_ 7.4, *p* value for pH_o_ 7.0 and 6.0 were 0.15 and 0.06, respectively. Background electrical activity before application of PS are subtracted in all quantitative analysis. (e) Time course of TRPM3 currents elicited by voltage ramps ranging −100 to +100 mV. PS was applied continuously while reducing extracellular pH without allowing any washing period between subsequent extracellular solution applications. (f) Representative recording of TRPM3 current in (e) by ramp protocols ranging from −100 to +100 mV at the indicated pH_o_. (g) Mean current amplitude of TRPM3 at the indicated pH_o_ in (E) (mean ± SEM; n = 7, * indicates p<0.05 by unpaired Student’s t-test; “n.s.” indicates not statistically significant). (h) Current amplitude at the indicated pH_o_ normalized to the current amplitude at pH_o_ 7.4.

**Figure 2.**
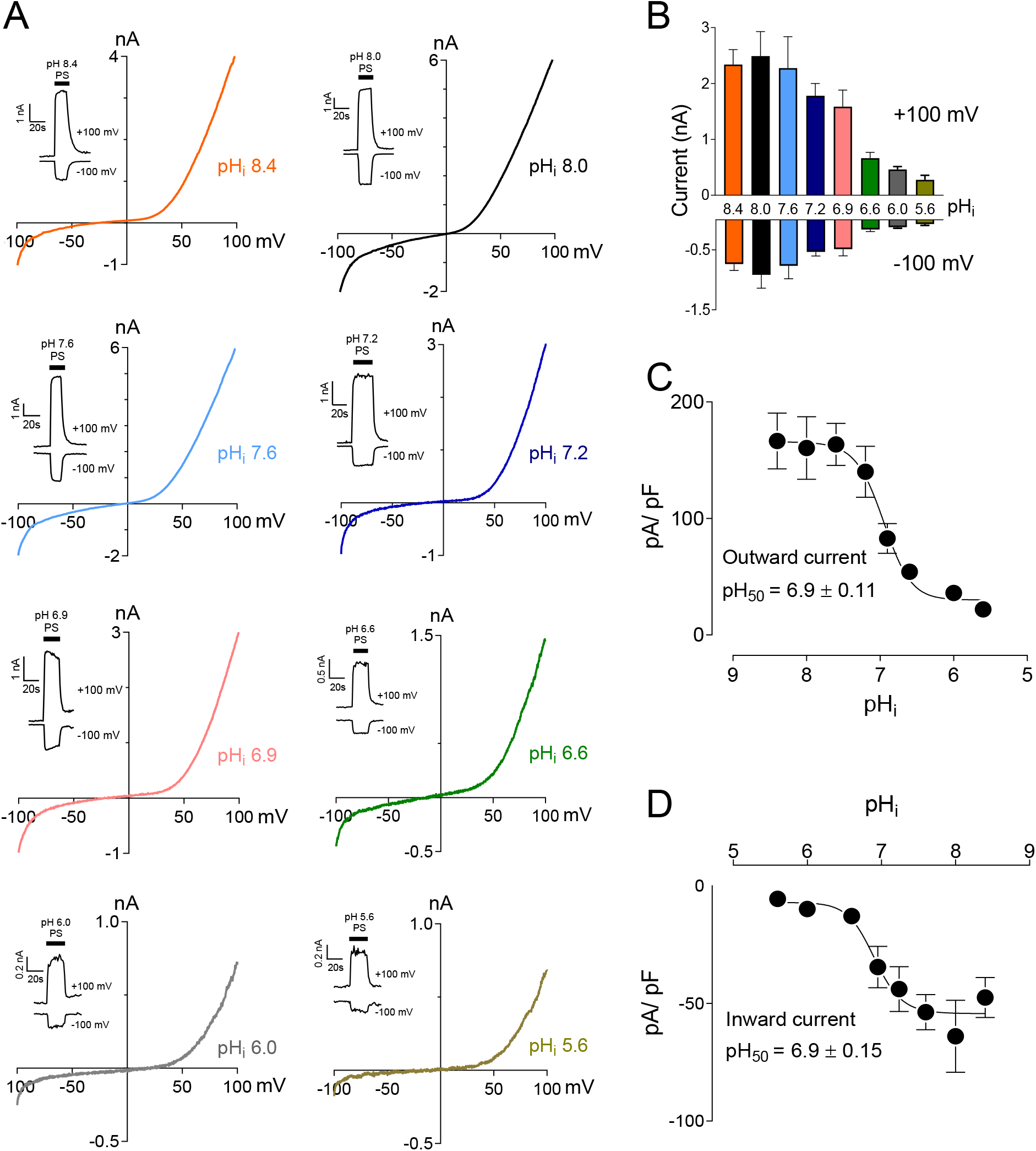
Intracellular acidification blocks TRPM3 activation by PS in a concentration-dependent manner. (a) Representative recordings and time courses (insert) of TRPM3 current by ramp protocols ranging from −100 to +100 mV at the indicated pH_i_. PS was applied through the extracellular solution (pH_o_ 7.4) and different cells were exposed to different pH_i_ while keeping that pH_i_ constant. (b) Mean current amplitude of TRPM3 at the indicated pH_i_ (mean ± SEM; n = 8 - 14). (c and d) pH_i_ concentration-dependence of TRPM3 activation by PS. c and d show outward and inward current, respectively, elicited by TRPM3 after extracellular PS application, by voltage ramp ranging −100 to +100 mV. TRPM3 currents exerted at −100 and +100 mV were considered as inward and outward current, respectively, and were utilized for these plots. All currents were normalized to corresponding capacitance of the cell overexpressing hTRPM3-GFP. Each data point is the mean of 8-14 cells with the error bar showing SEM, at the indicated pH_i_. Inhibitory pH_50_ values were measured separately for outward (pH_50_ = 6.9 ± 0.11) and inward (pH_50_ = 6.9 ± 0.15) currents.

### Effect of low pH_i_ on concentration-dependence of TRPM3 activation by PS

PS binds directly to the extracellular side of the TRPM3 channel to activate it. TRPM3 channel stays both functional and unaffected by the presence of intracellular PS (Wagner et al. 2008). Previous studies have shown that PS activates TRPM3 in a concentration-dependent manner with an EC_50_ value of 12 μM and 23 μM for outward and inward current respectively (Wagner et al. 2008). To investigate the effect of protons on PS concentration-dependence of TRPM3, we perfused the cells with a wide range of PS concentrations (1 μM - 500 μM), while keeping pH_i_ constant at 7.2 or 6.0. TRPM3 whole-cell outward and inward currents showed very similar PS concentration-dependence in both pH_i_ conditions (Fig. 3). EC_50_ values for outward currents were 16 μM (pH_i_ 7.2) and 15 μM (pH_i_ 6.0) (Fig. 3D), and for inward currents were 21 μM (pH_i_ 7.2) and 26 μM (pH_i_ 6.0) (Fig. 3F). All concentration-dependent curves plateaued after 50 μM PS stimulations (Fig. 3). We observed a downward shift of the outward current ratio curve in response to low pH_i_ (Fig. 3C). However, when currents were normalized to the maximum TRPM3 activation by 500 μM PS, the outward current curve did not deviate in response to low pH_i_ (Fig. 3D). These results indicate that higher proton concentration inside the cell reduces TRPM3 maximal activation potential at any given PS concentration but does not affect PS concentration-dependent activation of TRPM3. The PS concentration-dependent curve of TRPM3 inward currents did not show any change in response to low pH_i_ (Fig. 3, E and F). In summary, these data suggest that intracellular protons do not compete with PS for binding sites.

**Figure 3.**
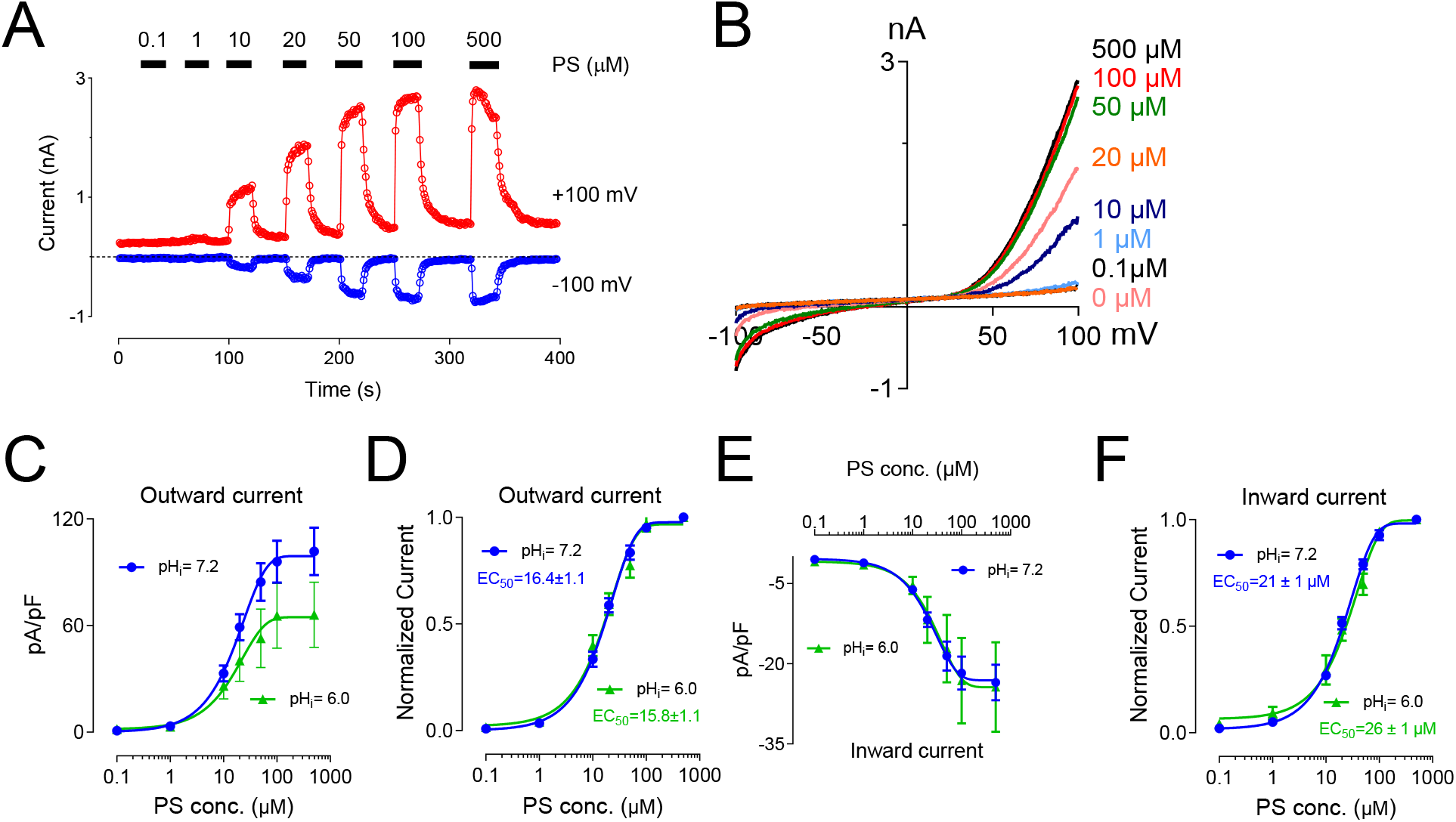
Concentration–response curve for PS-induced currents in hTRPM3, at the indicated intracellular pH. (a) Time course of TRPM3 currents elicited by voltage ramps ranging −100 to +100 mV. PS was applied extracellularly in increasing concentration sequence at concentrations of 0, 0.1, 1, 10, 20, 50, 100 and 500 μM, with adequate washing period between subsequent PS applications. (b) Representative recording of TRPM3 current at the indicated PS concentration in (a). (c) Mean outward currents measured at+100 mV and normalized to corresponding capacitance of the cell, error bars showing SEM (n = 7 cells), plotted against logarithmic values of PS concentrations. (d) Outward current normalized to maximum concentration-response (500 μM) of the same cell in (c). The EC_50_ values of PS for the outward currents are 16.4 ± 1.1 μM at pH 7.2 and 15.8 ± 1.1 μM at pH 6.0. (e). Mean inward currents measured at −100 mV and normalized to corresponding capacitance of the cell, error bars showing SEM (n = 7 cells), plotted against logarithmic values of PS concentrations. (f). Inward current normalized to maximum concentration-response (500 μM) of the same cell in (c). The EC_50_ values of PS for the inward currents are 21.0 ± 1.0 μM at pH 7.2 and 26.0 ± 1.0 μM at pH 6.0.

### TRPM3 inhibition by protons can be reversed by increasing pH_i_

Extracellular application of NH_4_Cl produces a rise in pH_i_ resulting from an influx of NH_3_. NH_3_ binds intracellular protons and causes alkalization inside the cells (Jacobs 1922; Warburg 1922). We observed that protons blocked TRPM3 from the cytoplasmic side, thus we tested whether raising pH_i_ by applying NH_4_Cl could rescue the TRPM3 current. We perfused the cells with solutions where part of NaCl was replaced by an equal amount of NH_4_Cl (see experimental procedures), to test whether increasing pH_i_ while keeping extracellular pH the same can reverse the blocking effects of protons on TRPM3. As shown in Fig. 4, A and B, 30mM NH_4_Cl increased recorded TRPM3 currents significantly. We attribute this increase to the influx of NH_3_ into the cell and removing bound protons from the TRPM3 cytoplasmic side. Overall, this result substantiates our finding that TRPM3 is blocked by protons binding to an intracellular site. Different concentrations of NH_4_Cl were applied to the same cell, and all NH_4_Cl applications change intracellular pH. To confirm that every TRPM3 stimulation starts at the same initial pH_i_, we allowed adequate washing time between two consecutive NH_4_Cl + PS applications.

**Figure 4.**
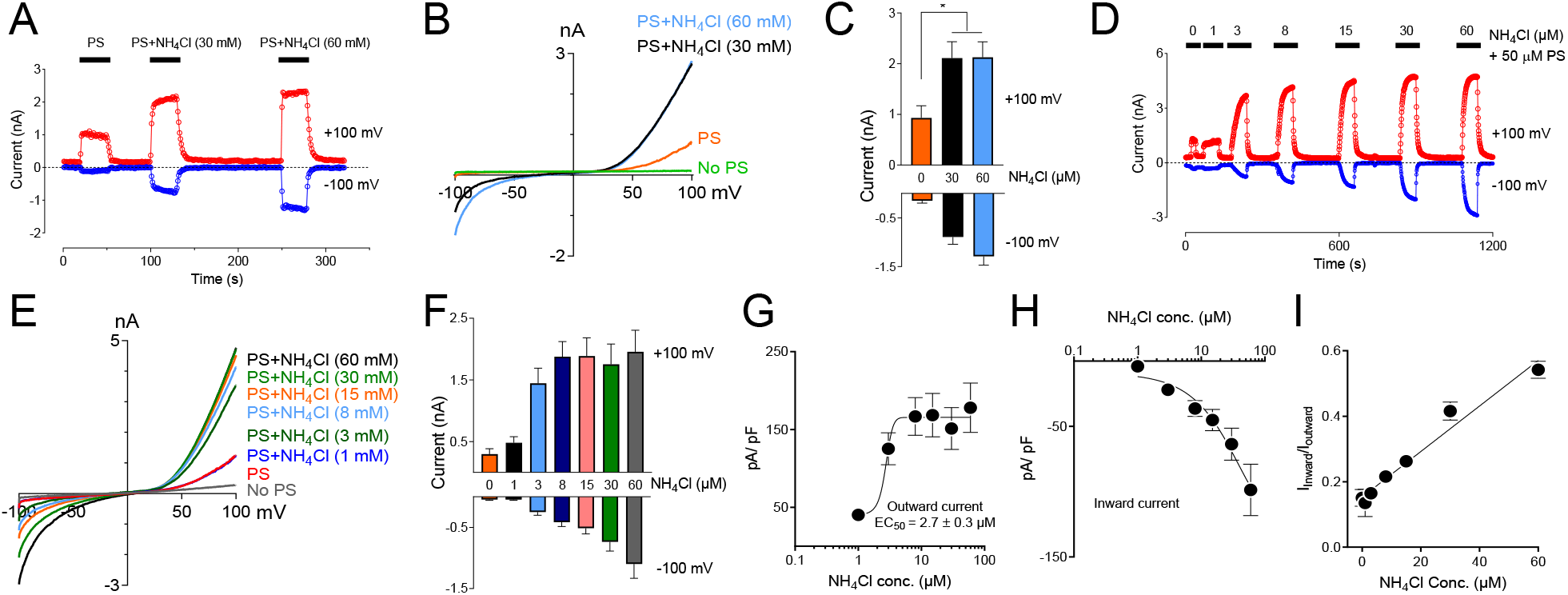
Inhibitory effect of low intracellular pH on TRPM3 can be reversed by perfusing cells with extracellular solution containing NH_4_Cl. (a) Time course showing outward and inward current, recorded at +100 and −100 mV respectively, obtained from HEK cells overexpressing TRPM3, under voltage ramp protocol ranging −100 to +100 mV, (pH_i_ = 6.0, n = 11 cells). Indicated concentrations of NH_4_Cl was applied extracellularly along with PS. To achieve similar osmolarity, all extracellular Na^+^ concentration was lowered to 85mM and osmolarities were adjusted to 300 ± 10 mOsm by mannitol. Adequate washing time (1 - 3 minutes) was provided after each application of NH_4_Cl, to bring the current down to the basal level, which is 6.0 for all the cells recorded. Representative current was plotted versus time in the presence of extracellular NH_4_Cl and PS, at the indicated concentrations. To achieve similar osmolarity, all extracellular buffer Na^+^ concentration was lowered to 85mM and osmolarities were adjusted to 300 ± 10 mOsm by mannitol. (b) Representative recording of TRPM3 current by ramp protocols ranging from −100 mV to +100 mV at the indicated NH_4_Cl concentrations. (c) Mean outward and inward TRPM3 current at the indicated NH_4_Cl concentrations (mean ± SEM, n=11 cells). * indicates p<0.05 by unpaired Student’s t-test. (d) Representative recording of TRPM3 current by the does-dependent effects of NH_4_Cl. The applications of extracellular NH_4_Cl at the indicated concentrations. (e) Representative recording of TRPM3 current by ramp protocols ranging from −100 mV to +100 mV at the indicated NH_4_Cl concentration. (From panel d) (f) Mean outward and inward TRPM3 current at the indicated NH_4_Cl concentration. (From replicated experiment of panel D) (mean ± SEM, n=11 cells). (g&h) Concentration-dependence curves of effects of NH_4_Cl on TRPM3 activations, outward (g) and inward (h) currents, respectively. All currents were normalized to corresponding capacitance of the cell overexpressing hTRPM3 (mean ± SEM, n=11 cells) at the indicated NH_4_Cl concentration. EC_50_ values were measured separately for outward (NH_4_Cl_50_ = 2.7 ± 0.3 μM) currents, whereas the inward currents continues to increase after applying 60 mM NH_4_Cl. (i) Mean ratio of inward current to outward current plotted against NH_4_Cl concentrations (mean ± SEM, n=11 cells).

Subsequently, we wanted to test whether external NH_4_Cl exhibits a concentration-dependent rescuing effect on TRPM3 activity or not. We perfused hTRPM3-GFP overexpressed HEK-293 cells with modified Tyrode solutions containing different concentrations of NH_4_Cl (1 - 60 mM). Na^+^ concentrations in all solutions were kept the same, and osmolarities were adjusted to 300 ± 10 mOSM with Mannitol. We observed a 5-fold increase in TRPM3 activity at only 3 mM (NH_4_Cl) concentration. Even 1 mM NH_4_Cl rescued some TRPM3 activity (p<0.05). NH_4_Cl concentrations higher than 3 mM had higher impact on TRPM3 activity than 3 mM NH_4_Cl. Although, when compared with 3 mM NH_4_Cl, some of these higher concentrations showed significant increase in TRPM3 activity, there were no substantial increase in TRPM3 activity by further increase in NH_4_Cl concentration. This data indicates that TRPM3 activity is highly sensitive to intracellular pH changes, at least at the intracellular pHs over 6.0. It is noteworthy that we did not measure exact pH_i_ changes following external NH_4_Cl applications, so the pH_i_ differences, before and after NH_4_Cl applications, are unknown here. However, these experiments still substantiate our claim that TRPM3 activity is highly sensitive to intracellular pH changes, at least at physiological pH_i_ ranges (pH_i_ 6.0 to 8.0). In this experiment, TRPM3 current recordings from each cell spanned over 20 minutes period. To confirm that subsequent application of NH_4_Cl and PS for a long time does not affect TRPM3 activity, we also tested individual cells with only one exposure to NH_4_Cl, at different concentrations. The results of these experiments are summarized in Supplementary Fig. 1 (sFig. 1). When compared with TRPM3 currents from our earlier experiments (Fig. 4, D), where the same cells were perfused with different concentrations of NH_4_Cl, individual cells did not show a difference (sFig. 1B). We also observed a higher increase in inward current than corresponding the outward current (Fig. 4, I). The ratio of inward current to outward current increased from 0.13 at 1mM NH_4_Cl to 0.54 at 60mM NH_4_Cl. This increase was consistent with the result obtained from individual cells (sFig. 1).

### TRPM3 is potentially permeable to protons

For a better understanding of the proton inhibition of TRPM3, we hypothesized that protons permeate through TRPM3 when it is activated by PS. The lipid bilayer (*e.g*. cell membrane) presents a strong barrier for the transport of charged ions through eukaryotic cell membranes. Although the permeability of protons is higher than other monovalent cations, which can partially be explained by the presence of transient water wires or long-lived hydrophilic pores (Tepper and Voth 2005), it is unlikely that these mechanisms can transport sufficient protons across the membrane to have a direct impact on an overexpressed protein, unless protons are passing through the overexpressed ion-channel itself. We thus examined whether protons can permeate through TRPM3. We recorded TRPM3 inward current in HEK-293 cells, overexpressing hTRPM3-GFP, by holding the membrane at −100mV and applying PS and low pH solutions from the outside. We maintained pH_i_ at 7.6 to provide a higher concentration gradient for protons. All cations except protons were removed using NMDG in external and internal solutions. PS, along with NMDG pH 5.5 extracellular solution, produced a small transient inward current in TRPM3 transfected HEK-293 cells (Fig. 5). Indeed, the amplitude of this current is significantly lower than TRPM3 currents observed in our whole-cell recordings. This is because in this experiment, all cations have been removed besides a limited amount of H^+^. extracellular and intracellular proton concentrations was 3.2 × 10^-3^ mM (pH 5.5) and 2.5 ×10^-5^ mM (pH 7.6) respectively. These concentrations represent the total cation concentrations of these solutions, which are several orders of magnitude lower than the total cation concentrations of our modified Tyrode solutions. In addition, extracellular pH 5.5 blocks about 60% of TRPM3 currents. However, to provide a reasonable proton gradient across the cell membrane, it is critical to conduct this experiment at a low extracellular pH. Since, there were no other cation involved, although low in amplitude, this current suggests passage of proton through TRPM3 channel. In addition, the proton current is transient is because of the intracellular inhibitory effects of H^+^ on TRPM3 after its permeation. Mock transfected cells did not produce any inward current in response to PS (Data not shown). This evidence indicates that TRPM3 is potentially permeable to proton.

**Figure 5.**
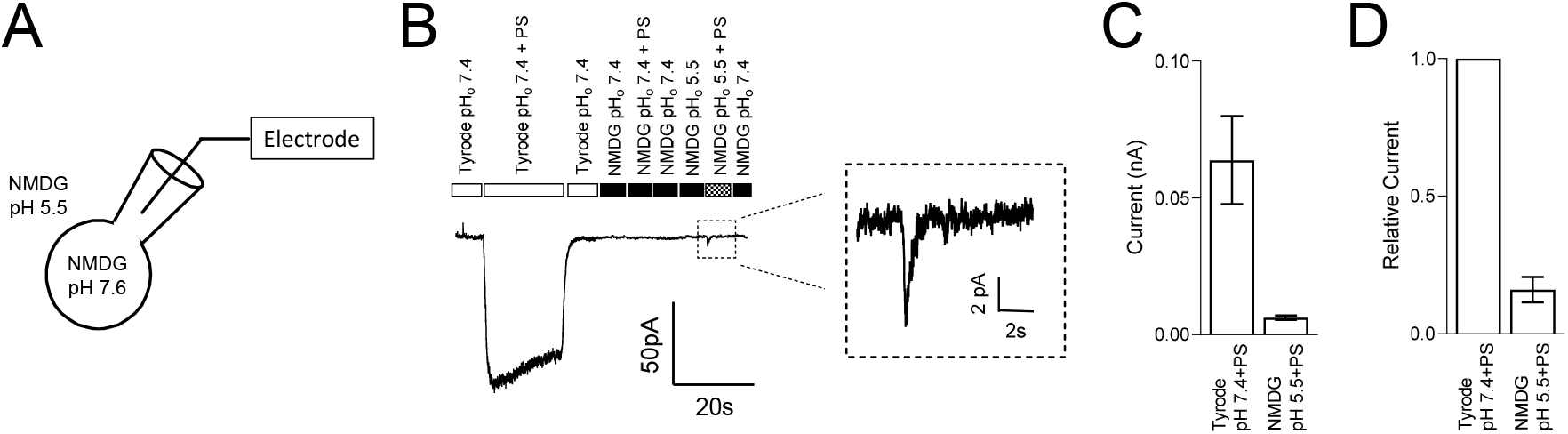
TRPM3 is potentially permeable to protons. (a) Schematic showing the protons permeation recording condition, where all intracellular and extracellular ions were replaced by NMDG and glutamic acid, respectively, except for protons. 50 μM PS was applied while holding hTRPM3-GFP transfected HEK cells at −100 mV. (b) Inward currents were elicited by PS application. No PS activated current under the NMDG solution at pH_o_ 7.40, while lowering down the pH_o_ to 5.50 generated a small and transient inward current. Insert panel shows the inward current recorded during NMDG-PS (pH 5.5) application. (c) Mean current amplitude in response to PS at the indicated conditions (mean ± SEM, n = 8 cells). (d) Relative current amplitude in response to PS, comparing with Tyrode (pH_o_ 7.4) solution response of the same cell (mean ± SEM, n = 8 cells).

### The inhibition of intracellular Ca^2+^ on TRPM3 is pH independent

TRPM3 displays higher permeability for divalent cations than monovalent cations. For the splice variant TRPM3α2, 24% of total TRPM3 current is expected to result from Ca^2+^ (Przibilla et al. 2018), suggesting a large increase in [Ca^2+^]_i_ following TRPM3 activation. Multiple studies verified this effect showing an increase in [Ca^2+^]_i_ in a micromolar range following TRPM3 activation (Vriens et al. 2011; Straub et al. 2013; Przibilla et al. 2018). A recent study demonstrated that an increase in [Ca^2+^]_i_, independent of TRPM3 activity, inhibits TRPM3 in a calmodulin-dependent manner (Przibilla et al. 2018). This suggests a potential negative feedback mechanism that regulates a high increase in [Ca^2+^]_i_ resulting from TRPM3 activation. Studies conducted by others enlist [Ca^2+^]_i_ as a regulator of TRPM3 activity. In our study, we found that intracellular proton blocks TRPM3 as well. Hence, we asked the question, how do these two regulatory mechanisms interact with each other? To find the effect of [Ca^2+^]_i_ on the concentration-dependent inhibition of TRPM3 by protons, we tested TRPM3 activity in two different [Ca^2+^]_i_ conditions, while providing a wide range of pH_i_, using whole-cell patch clamp of TRPM3 transfected cells. In addition to the inhibition of TRPM3 current by [Ca^2+^]_i_ observed by J. Przibilla et al., we observed an additional delayed inhibition of TRPM3 current by Ca^2+^. For example, under 1μM [Ca^2+^]_i_, recorded current showed a further inhibition after initial activation and the residual current amplitudes were less than 50% of the initial activation (Fig 6, B and C). Low [Ca^2+^]_i_ (< 10 nM) did not show a delayed inhibition of TRPM3 current (Fig. 6A). pH_i_ concentration-dependence of TRPM3 was not affected by [Ca^2+^]_i_, as they showed similar pH_50_ Values (7.0 ± 0.1, 7.1 ± 0.1, 7.2 ± 0.5) for low [Ca^2+^]_i_, high Ca^2+^ initial and high Ca^2+^ delayed current intensities (Fig. 6E). It is noteworthy that, despite having similar pH_50_ values, TRPM3 outward current densities were significantly decreased in high [Ca^2+^]_i_ conditions (Fig. 6E). To investigate if protons affected the percentage of current inhibited in delayed current intensities compared with the initial current intensities, we analyzed the ratio of delayed to initial current intensities in different pH_i_ at high [Ca^2+^]_i_ concentration. We did not observe any significant difference between the ratios resulting from different pH_i_ (Fig. 6F). Overall, these results indicate that, although [Ca^2+^]_i_ inhibits TRPM3, it does not affect the regulation of TRPM3 by intracellular protons.

**Figure 6.**
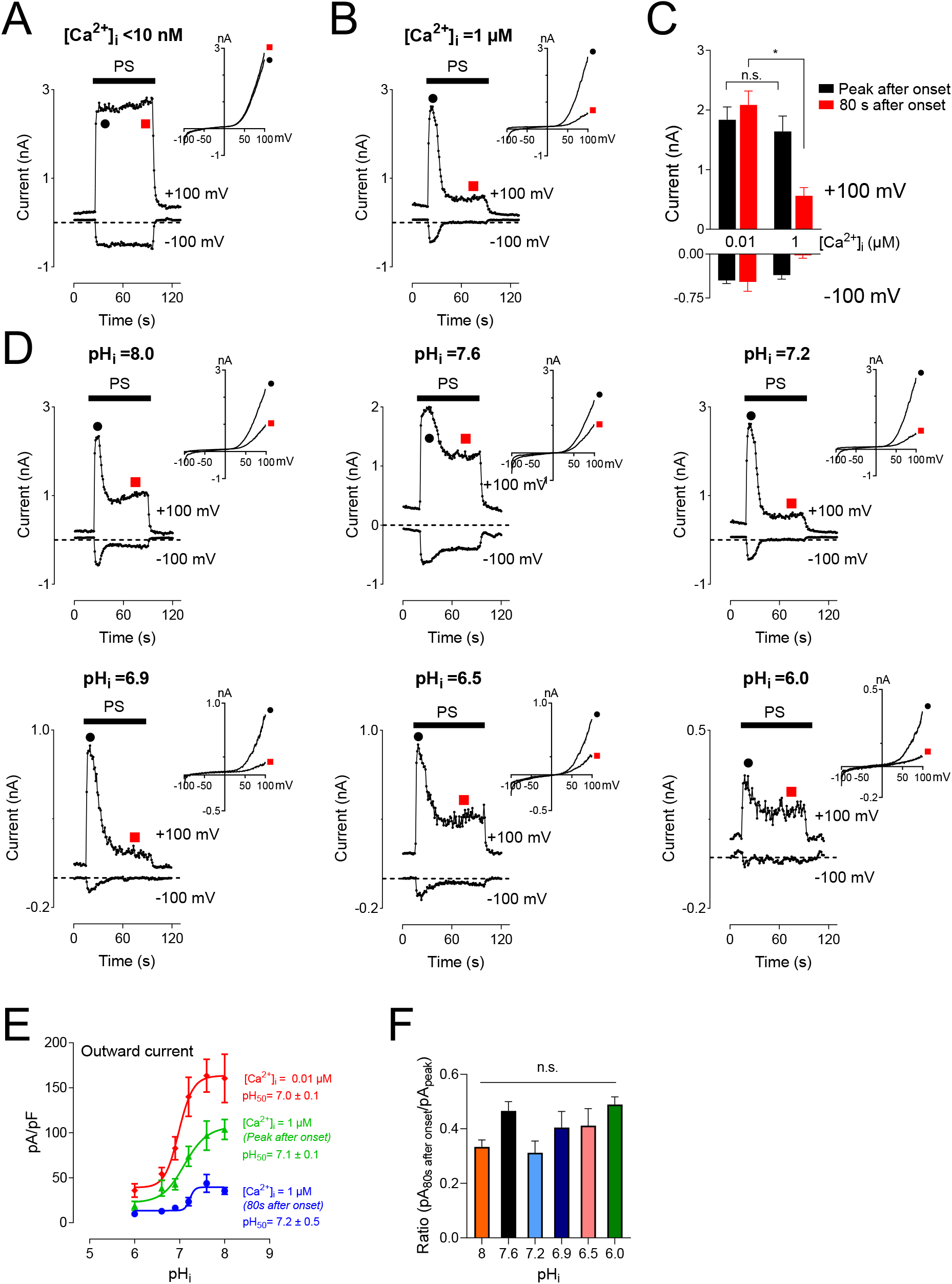
Inhibitory effect of intracellular high Ca^2+^ on TRPM3 activation by PS. (a and b) Time course of TRPM3 currents elicited by voltage ramps ranging −100 to +100 mV at the indicated [Ca^2+^]_i_ concentration. At high [Ca^2+^]_i_ (1 μM), TRPM3 is activated by PS initially (Black symbol), but runs down soon afterwards (Red symbol). Whereas the TRPM3 current was sustained at the low [Ca^2+^]_i_ condition. Inserts show the representative recordings from the indicated points. (c) Mean current amplitude of TRPM3 at the indicated [Ca^2+^]_i_ (mean ± SEM; n = 10). * indicates p<0.05 by unpaired Student’s t-test; “n.s.” indicates not statistically significant. (d) Time course and representative recordings of TRPM3 current with [Ca^2+^]_i_ (1 μM) at the indicated pH_i_ from pH_i_ 8.0 to pH_i_ 6.0. (e) pH concentration-dependence of TRPM3 activation by PS at the indicated [Ca^2+^]_i_. Green and blue symbols both represent high [Ca^2+^]_i_, while green represents currents elicited right after PS application (peak after onset), and blue represent currents remaining after inhibition of by high [Ca^2+^]_i_ (80s after onset). All currents were normalized to corresponding capacitance of the cell overexpressing hTRPM3. Each data point is the mean of 8-14 cells with the error bar showing SEM, at the indicated pH_i_. (f) Mean ratio of the peak current after onset to the current 80 seconds after onset (mean ± SEM, n = 8-14 cells). “n.s.” indicates not statistically significant among groups by one-way ANOVA with Tukey’s post hoc multiple comparison.

### Molecular mechanism underlying TRPM3 inhibition by protons

Glutamate, aspartate, histidine, and lysine residues are potential proton acceptors, especially glutamate and aspartate, which present negative charges (Zhou and Pang 2018). To identify the amino acid residues of TRPM3 accountable for its sensitivity to pH_i_, we prepared hTRPM3-GFP mutant plasmids having mutations in the pore region. We selected all eight glutamate and aspartate amino acids, in the vestibule of the loop between S5 and S6 transmembrane domains (Fig. 7A). These eight residues were either a glutamate and aspartate amino acid, and they were mutated to glutamine. We created two double mutants (E1034Q - E1035Q, E1072Q - D1073Q) and six single mutants (E1055Q, D1059Q, D1062Q, E1069Q, E1072Q, and D1073Q). We expressed these mutants in HEK-293 cells and recorded elicited currents in response to PS, while perfusing with physiological and low pH external solutions (Fig. 7B). Except for E1055Q, which resulted in a non-functional ion-channel, all other mutants exhibited identical I-V relation to WT-hTRPM3, although most of the mutants showed markedly reduced current amplitudes (Fig. 7B). We perfused cells expressing these mutants with external solutions of pH 7.4 and pH 5.5. The pH of 5.5 was selected as representative of low pH external solutions because at this pH, WT-TRPM3 currents were blocked significantly yet sufficient activity was maintained for analysis. We compared the percent decrease in TRPM3 current from pH_o_ 7.4 to pH_o_ 5.5 for all the mutants and WT-TRPM3 (Fig. 7C). The double mutant (E1034Q - E1035Q), E1069Q, and E1072Q showed similar sensitivity to protons when compared with the WT-TRPM3.

**Figure 7.**
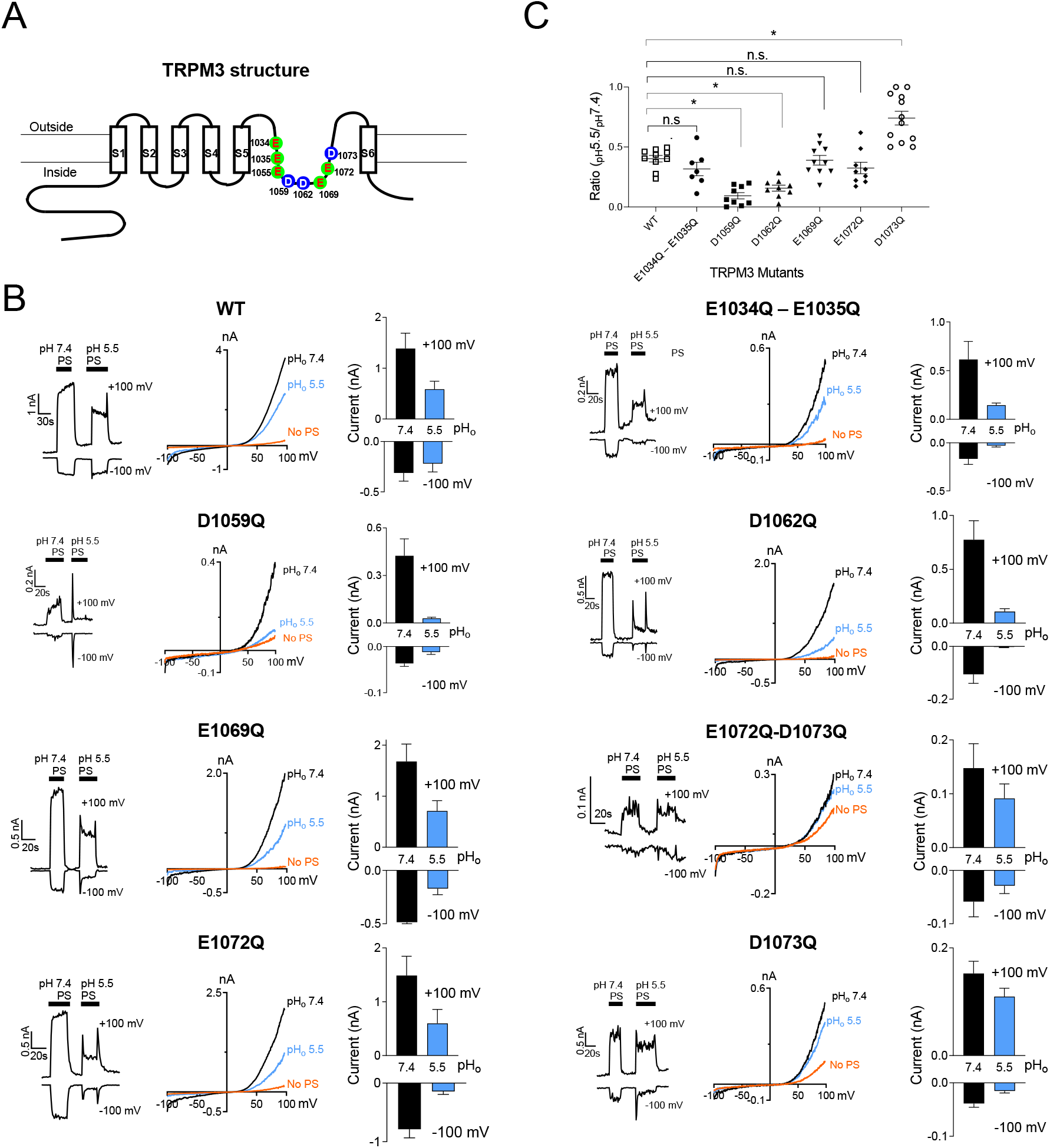
Changes of protons sensitivity of TRPM3 mutants. (a) Schematic of TRPM3 structure and the substituted amino acid residues in the putative pore region of hTRPM3. (b) Time course and representative recordings of TRPM3 mutants and wild-type currents elicited by voltage ramps ranging −100 to +100 mV, at the indicated pH_o_. Bar graphs show mean outward and inward current amplitudes at the indicated pH_o_ (mean ± SEM, n = 7-12 cells) Internal solution had a constant pH of 7.20 for all the recordings. (c) Ratio of outward current amplitudes at pH_o_ 5.5 and pH_o_ 7.4, of TRPM3 mutants and WT control. To obtain these ratios, current elicited by +100mV at pH_o_ 5.5 was divided by the current elicited by +100mV at pH_o_ 7.4 of the same cell (n = 7-12 cells). P values, comparing each group with WT-TRPM3, * indicates p<0.05 by unpaired Student’s t-test; “n.s.” indicates not statistically significant.

Mutants D1059Q and D1062Q were found to be more sensitive to protons, as the reduction of whole-cell current due to low pH_o_ was increased to 93.9% and 84.1% in D1059Q and D1062Q, respectively, compared to 56.5% observed in WT-TRPM3. Mutant D1073Q showed significantly less sensitivity towards protons, as pH_o_ 5.5 reduced its current amplitude only by 25%. As summarized in Fig. 7C, these results establish the amino acid residues D1059Q, D1062Q and D1073Q as some of the key determinants of protons sensitivity in TRPM3. Although it is unclear at this moment why mutating these residues produce variable effects (increased or decreased proton sensitivity), these data suggested that the pore vestibule of TRPM3 is critical for the pH sensitivity. Further studies are required to delineate the underlying mechanism of these variable effects.

## Discussion

We demonstrated for the first time that TRPM3 is a proton-permeable channel regulated by both extracellular and intracellular acidic conditions. Our experiments suggest a direct interaction of intracellular protons with the cytoplasmic side of TRPM3 to induce a blocking effect, whereas extracellular protons permeate through the ion-channel first to block it from the intracellular side. We also demonstrated that the blocking effect of the intracellular protons could be reversed by decreasing intracellular protons concentrations, which indicates a reversible binding of protons to the intracellular amino acid residues. We identified an internal residue responsible for protons sensitivity of TRPM3. Finally, we demonstrated that low internal pH produces a downward shift of PS concentration-dependent activation of TRPM3 by reducing the amplitude of TRPM3-mediated currents at any given pH_o_. Overall, we report evidence for the regulatory role of protons on TRPM3 activation and the molecular mechanism responsible for it.

### General characteristics of TRPM3

TRPM3 is a heat-sensitive ion channel, which is expressed in a somatosensory neuron where its role as a noxious heat sensor has been established (Vriens et al. 2011; Vandewauw et al. 2018). Activation of pancreatic TRPM3 increases glucose-induced insulin release (Wagner et al. 2008). These data indicate the existence of multiple TRPM3 regulatory molecules. Indeed, cytosolic phosphatidylinositol bisphosphates (PIPs) and ATP have a stimulatory effect on TRPM3. Activation of phospholipase C-coupled muscarinic acetylcholine receptors inhibit recombinant and endogenous TRPM3 (Toth et al. 2015). Calmodulin binds to multiple sites of TRPM3 in Ca^2+^-dependent manner, and both intracellular calmodulin and Ca^2+^ inhibits TRPM3 (Holakovska et al. 2012; Przibilla et al. 2018). Here we report the modulatory role of protons on TRPM3. Reversible inhibition of TRPM3 by protons indicates that acidic pH may serve as a negative feedback mechanism to regulate TRPM3 activity in physiological/pathological conditions.

### Regulations of pH on TRP channels

Acidification has modulatory effects on variety of ion channels (discussed in the introduction) including TRP channels. Extracellular acidic pH modulates TRPV1 channel gating (Jordt, Tominaga, and Julius 2000; Ryu, Liu, and Qin 2003), stimulates TRPC4 and TRPC5, but inhibits TRPC6 (Semtner et al. 2007), TRPM5 (Liu, Zhang, and Liman 2005), and TRPV5 (Yeh et al. 2003), and potentiates TRPM6 and TRPM7 inward currents (Jiang, Li, and Yue 2005; Li, Jiang, and Yue 2006; Li et al. 2007). Intracellular protons inhibit TRPM7 (Kozak et al. 2005), and block TRPM8 (Andersson, Chase, and Bevan 2004). Here we demonstrate that PS-induced TRPM3 outward and inward currents both are inhibited by protons. Although both intracellular and extracellular acidic conditions inhibit TRPM3, there are differences between internal and external proton-induced inhibition. Extracellular protons cannot inhibit TRPM3 until pH reaches below 6.0, but pH_i_ has a pH_50_ value of 6.9. We demonstrated a very sharp inhibition by protons at an external pH below 6.0; this inhibition did not follow a typical concentration-response relationship. Whereas internal protons efficiently blocked TRPM3 current and showed a concentration-response relationship. These results suggest that protons are not competing with PS for the same binding site of TRPM3, and indicated that TRPM3 may contain proton-binding sites in the cytoplasmic domains. It is plausible that an increase in extracellular proton’s concentration causes protons influx through TRPM3 that enables protons to bind to its internal binding site to inhibit TRPM3. To rule out the effects of proton-activated endogenous chloride currents, we conducted all whole-cell recordings under very low intracellular and extracellular Cl^-^ concentrations. Indeed, our external and internal solutions contained 8 mM Cl^-^, but this concentration of protons is not capable of activating endogenous anions channels of HEK293 cells (Lambert and Oberwinkler 2005).

### Mechanisms of effects of protons on TRPM3

It appeared that extracellular pH (< 6.0) strongly blocks TRPM3 activities, which would indicate TRPM3 is regulated by extracellular acidic pH directly. However, the lack of pH concentration-dependent activity did not support this hypothesis. These findings led us to the conclusion that protons do not have extracellular binding sites, at least under the activations of PS. Reduced intracellular pH appeared to have clear pH concentration-dependent effects on TRPM3, which begs the question: how might external protons affect TRPM3 gating properties? The cell membrane must have a mechanism for the proton permeation and thus change the intracellular pH. It is possible that protons cross the TRPM3 while it is open, although we cannot exclude the possibility that protons may permeate the cell membrane directly.

Our data suggests that protons directly cross TRPM3 when it is activated by PS. Consistently, no proton current was found without activating TRPM3, supporting the conclusion that TRPM3 is potentially permeable to protons. We also studied the regulation of TRPM3 by Ca^2+^. Previous studies have shown that TRPM3 is a Ca^2+^ permeable ion channel (Grimm et al. 2003; Lee et al. 2003). TRPM3 channel activity strongly depends on intracellular Ca^2+^ (Przibilla et al. 2018). Along with the inhibition of TRPM3 current amplitude, we also found that Ca^2+^ accelerates the decay time of TRPM3. Increased [Ca^2+^]_i_ from minimum to 1μM significantly reduces the plateau of the current. In general, [Ca^2+^]_i_ is a key regulator to modulate the TRPM3 gating - we hoped to visualize the interaction between [Ca^2+^]_i_ and protons. However, high [Ca^2+^]_i_ did not change the pH_50_ of the effects of intracellular protons on TRPM3 (Fig. 6). Vice versa, decreased [H^+^]_i_ did not affect the Ca^2+^ inhibition on TRPM3 (Fig. 6). Therefore, it appears unlikely that protons can inhibit TRPM3 channel activities by competing with [Ca^2+^]_i_ for a binding site on the cytoplasmic side.

To determine the molecular mechanism by which intracellular protons inhibit TRPM3, we mutated all the titratable residues in the pore region between S5 and S6 to determine which sites are responsible for pH_i_ sensitivity. Among most of the mutations of Asp and Glu residues, three residues-D1059, D1062 and D1073-which we predicted to locate in the inner vestibule in the pore region, strongly change the proton sensitivities. We conclude these residues in the pore region could be the proton binding sites. Of course, we should not exclude any other intracellular binding sites. For example, C terminus of the S4-S5 linker is thought to be critical for changing TRP channel’s pH_i_ sensitivity (Du, Xie, and Yue 2009). Although, how intracellular protons change TRPM3 gating properties through these residues is still unknown, it will be of interest to investigate whether acidic intracellular pH alters intracellular signaling pathways in future studies. It will also be of interest to investigate other potential proton binding sites that might interact with [Ca^2+^]_i_ near the intracellular mouth and act as the [Ca^2+^]_i_-activating site to regulate [Ca^2+^]_i_ mediated TRPM3 activation. Further investigation is required to test this hypothesis.

### Conclusion

Our findings suggest that cellular acidification serves as a negative or protective feedback mechanism to limit TRPM3 activities. Although the development of intracellular acidosis has not been well established, metabolic acidosis is a relatively common condition that causes pH_i_ to fall (Salameh, Ruffin, and Boron 2014). Some reports suggest that during metabolic acidosis, insulin secretion is depressed (Mak 1998; Bigner et al. 1996). Thus, it is possible that a low pH-mediated dampening of TRPM3 activity might contribute to the decreased insulin secretion observed in metabolic acidosis. If true, then modulating TRPM3 activity might be a potential future clinical application in treating acidosis induced pancreatic disorders.

Collectively, we show that external and internal acidic pH show strong and state-dependent inhibition of the TRPM3 channels. Asp1073 residue in the inner vestibule of the channel pore is critical in modulating this inhibition. Given the physiological significance of TRPM3 in numerous cells, including pancreatic beta cells and sensory neurons, understanding TRPM3 gating by protons may generate new physiological and/or pathological insights.

## Acknowledgement

We thank Dr. C. Harteneck for providing the TRPM3 construct. We thank Dr. Lixia Yue for her suggestions and comments. We thank Dr. Boren Lin, Erin Koffman, Tyler Ortyl, Farzaneh Naghavi, and Kritika Singh for their comments. J.D. is supported by the National Institutes of Mental Health (5R01 MH113986). LRR is supported by the National Institutes of Health (R00HL119560, OT2OD023854, OT2OD026582)

## Author Contributions

J.D. conceived and supervised the project. J.D., MZ.HS. and L.X. designed the experiments with input from Y.S.L. and L.R.R. MZ.HS. and L.X. did most of the patch-clamp experiments. Y.S.L. and L.R.R. performed molecular biology experiments and oversaw the mutagenesis of TRPM3. MZ.HS. and J.D. drafted the manuscript with input from all authors contributed to finalizing the manuscript.

## Abbreviations

TRP: Transient Receptor Potential;
TRPM: Melastatin-like TRP channels;
GFP: Green Fluorescent Protein;
NMDG: N-methyl-D-glucamine;
PS: Pregnenolone Sulfate;
pH_o_: Extracellular pH;
pH_i_: Intracellular pH;
AP-1: Activator Protein-1;
Egr-1: Early Growth Response protein 1;
CRE: cAMP response element;
K2p: Two-pore domain K^+^ channels:
TWIK: Tandem of pore domains in Weak Inward rectifier K^+^ channels;
TASK: TWIK-related Acid-Sensitive K^+^ channels;
Kir: Inward rectifier K^+^ channel;
TREK: TWIK-RElated K^+^ channels;
TRESK: TWIK-RElated spinal cord K^+^ channels;
TALK: TWIK-related ALkaline pH-activated K^+^ channels;
ATP: Adenosine Triphosphate;
PKD2L1: Polycystic Kidney Disease 2-Like ion channel-1;
CGRP: Calcitonin Gene-Related Protein.

---

**Supplementary Figure 1.**
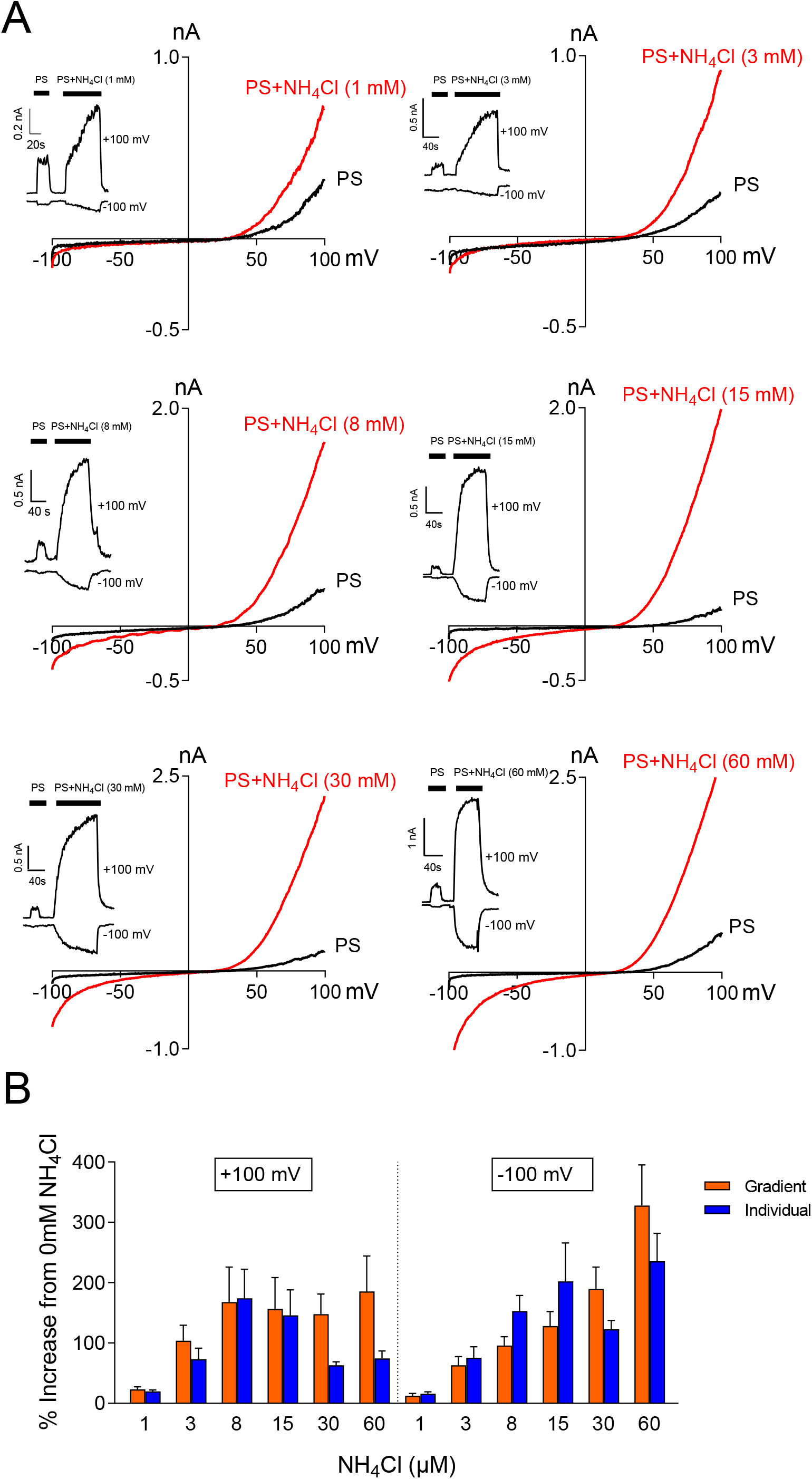
Concentration-dependent effects of NH_4_Cl on TRPM3 current. (a)Time course and representative recordings of TRPM3 current by ramp protocols ranging from −100 mV to +100 mV at the indicated NH_4_Cl concentrations. The pH_i_ was 6.0 in all recordings. Individual transfected cells were exposed to NH_4_Cl only once. (b)The comparison of effects of NH_4_Cl on TRPM3 currents in “gradient” (Fig. 4) and “individual” (a) recordings. The increase in outward current in response to NH_4_Cl is presented as the percentage increase in outward current from the same cell without NH_4_Cl. Both data suggest that NH_4_Cl potentiate both TRPM3 inward and outward currents. The average data are mean ± SEM, n = 11-20 cells.

